# Efficient generation of human oligodendrocyte grafts that restore myelin in adult human brain tissue

**DOI:** 10.1101/2025.02.28.640804

**Authors:** Raquel Martinez-Curiel, Oleg Tsupykov, Paula Rincón-Cerrada, Anna-Maissoun Al-Khani, Yogita Sharma, David Martín-Hernández, Emanuela Monni, Anita Adami, Ekaterina Savchenko, Laura R. Rodríguez, Srisaiyini Kidnapillai, Andreas Bruzelius, Isabel Hidalgo, Isaac Canals, Galyna Skibo, Daniella Rylander Ottosson, Anna Falk, Johan Bengzon, Henrik Ahlenius, Johan Jakobsson, Olle Lindvall, Zaal Kokaia, Sara Palma-Tortosa

## Abstract

Intracerebral transplantation of stem cell-derived oligodendrocytes (OLs) is a promising strategy for repairing demyelinated human brain tissue, the main hallmark of white-matter disorders. However, several challenges hinder clinical translation, including slow or inefficient production of human OLs with current protocols, and difficulty in generating pure OL grafts capable of remyelinating injured neural circuits. Here, we present a robust, highly reproducible method for the rapid and efficient production of human OLs from human induced pluripotent stem cell derived long-term neuroepithelial-like stem (lt-NES) cells. Induced expression of the lineage-defining transcription factors SOX10 and OLIG2 in lt-NES cells is sufficient to generate a population of 80% OLs within 7 days. Importantly, these cells survive, differentiate and form functional OL-exclusive grafts when transplanted into adult human brain slices *ex vivo*, constituting the first demonstration that an OL-exclusive graft with robust myelination capacity can be generated in a clinically relevant allogeneic environment. This advance marks a significant step towards the clinical application of oligodendrocyte replacement therapy for human demyelinating disorders.

## INTRODUCTION

Oligodendrocytes (OLs), the myelinating cells of the central nervous system (CNS), are essential for enhancing the propagation of action potentials, ensuring proper neuronal function. Their loss in white-matter disorders leads to axonal demyelination with critical neurological consequences. Current treatments for demyelinating disorders, such as leukodystrophies or multiple sclerosis, are only symptomatic, slowing down disease progression but not halting or reversing the myelin damage^1, 2, 3^. For this reason, novel therapies targeting remyelination processes are highly warranted. Importantly, in animal models, intracerebral transplantation of neural progenitor cells (NPCs) or already OL-committed progenitor cells derived from human pluripotent stem cells has shown potential for rebuilding the demyelinated CNS, improving motor and cognitive deficits^4–6^. The recovery has been attributed not only to the so-called bystander effect, i.e. by enhancing endogenous oligodendrogenesis, but also to actual OL replacement, as evidenced by graft-derived remyelination. However, from a clinical perspective, several challenges remain: **i**) Current protocols for generating human OLs for transplantation are time-consuming or inefficient in terms of yield^7–12^; **ii**) Due to limited access to adult human brain tissue, testing the capacity of generated OLs to myelinate human axons has been mainly restricted to *in vitro* co-cultures with either human induced pluripotent stem (iPS) cell-derived neurons or human fetal cortical neurons^7, 10, 11^; and **iii**) The generation of a pure OL population for transplantation, without off-target cell types, has yet not been achieved^9, 10^. In this context, the presence of astrocytes or neurons within the graft may impair remyelination in white matter lesions by contributing to glial scar formation or disrupting local neural circuits.

Here, we have developed and extensively characterized a fast and efficient protocol for generating a transplantable population of human OLs able to myelinate human brain tissue. By combining human iPS cell-derived long-term neuroepithelial-like stem (lt-NES) cells, which have previously been shown to produce a mixture of functional neurons and myelinating OLs^5^ after 21 days of cortical priming, with reprogramming technologies based on OLIG2 and SOX10 overexpression, we have achieved the generation of about 80% of human OLs after only 7 days of induction. Importantly, *ex vivo* transplantation of these cells into a clinically relevant human brain platform, adult human cortical organotypic slices, gives rise to OL-exclusive grafts, capable of myelinating host axons.

## RESULTS

### Overexpression of OLIG2 and SOX10 in human lt-NES cells efficiently induces OLs *in vitro*

Human iPS cell-derived lt-NES cells are long-term expandable, genetically stable and non-tumorigenic cells, which have shown therapeutic potential in several studies^5, 13–16^. We have previously shown that, following a cortical priming protocol, human lt-NES cells give rise to a mixture of mature neurons and functional OLs *in vitro* and after grafting into stroke-injured rat cortex and human organotypic cortical tissue^5, 15^. Myelinating OLs appeared in culture after 21 days of differentiation. However, only 5% of the cells acquired this fate^5^. To improve the yield of OLs, we tested alternative induction protocols based on the overexpression of OLIG2 and SOX10, transcription factors implicated in OL specification, using an inducible Tet-ON system combined with antibiotic selection to ensure co-overexpression in all cultured cells (**Fig. 1a**). Conversion efficiency into human induced OLs (hiOLs) was assessed by flow cytometry through quantification of O4+ cells, a marker expressed in OLs, at day 7 of induction. In addition, expression of markers for different neural cell populations was assessed by immunocytochemistry at day 10.

**Fig 1.**
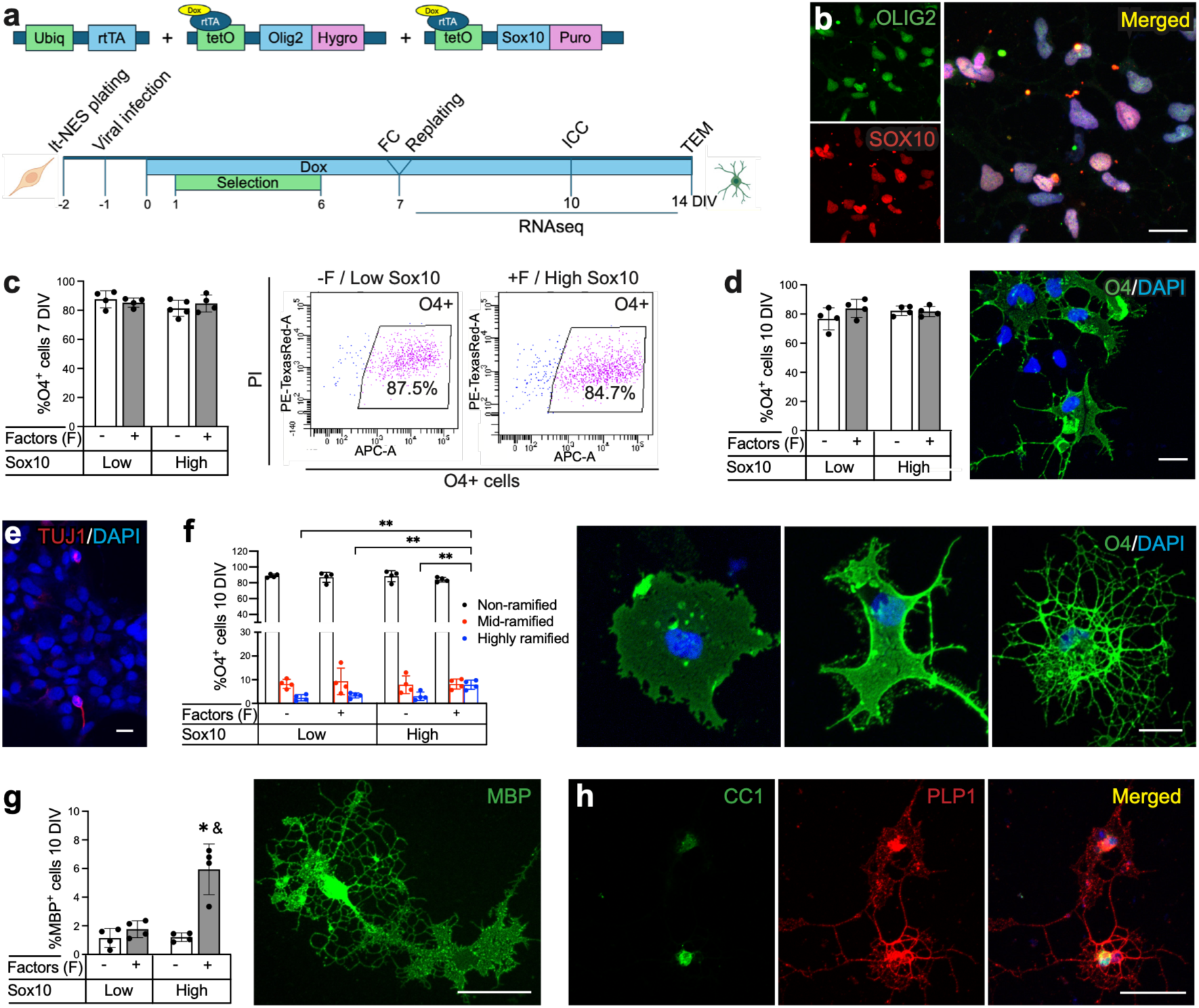
OLIG2 and SOX10 overexpression in human long-term neuroepithelial-like stem (lt-NES) cells efficiently generates oligodendrocytes (OLs) *in vitro*. **a** An experimental design for the production of human induced OLs from human lt-NES cells. FC–Flow cytometry, ICC–Immunocytochemistry, TEM–Transmission electron microscopy, Ubiq–ubiquitin, rtTA – reverse tetracycline-controlled transactivator, Puro–puromycin, Hygro– hygromycin and, Dox – doxycycline. Panel created in PowerPoint Version 16.107.4. **b** Confocal images showing expression of OLIG2 and SOX10 at day 10 of programming. Scale bar, 20 μm. **c** Quantification and two representative dot plots showing the percentage of the O4+ cells, OL marker, at day 7 using different programming protocols. **d** Quantification and representative confocal image of O4+ cells at day 10. Scale bar, 20 μm. **e** A confocal image showing expression of the neuronal marker TUJ1. Scale bar, 20 μm**. f-g** Confocal images and quantification of non-ramified, mid-ramified and highly ramified O4+ cells (**f**), and myelin basic protein (MBP)+ cells (**g**). Scale bar in **f**, 20 μm. Scale bar in **g**, 40 μm. **h** Representative images showing the co-expression of the mature OL marker CC1 and the proteolipid protein 1 (PLP1) in programmed cells at day 10. Scale bar, 40 μm. Nuclear staining (DAPI, blue) is included in the merged panels. All data are presented as the mean ± SD, n=4-5 biologically independent samples. Two-way ANOVA followed by Sidák’s multiple comparisons post hoc test. ** p<0.01, * p<0.05 vs. – factors + Low Sox10, & p<0.05 vs. – factors High Sox10.

We first attempted to produce OLs by overexpression of OLIG2 alone. However, the generated cells displayed neuronal morphology (**Supplementary Fig. 1a**) and expressed the neuronal markers doublecortin (DCX), beta-tubulin III (TUJ1), and microtubule-associated protein 2 (MAP2) (**Supplementary Fig. 1f**). Quantifications showed that 5.3 ± 3.8% of the cells expressed the OL marker O4, while 96.4 ± 1.4% of the generated cells were positive for the neuronal marker TUJ1 (**Supplementary Fig. 1c, e, h**). On the other hand, sole overexpression of SOX10, a protocol previously used on human iPS cells-derived NPCs to generate OLs^10^, gave rise to 48.3 ± 14.8% of O4+ cells within 7 days, while 41.9 ± 1.6% of the cells expressed TUJ1 and displayed typical neuronal morphology (**Supplementary Fig. 1b, d, e, g, h**). No expression of the astrocyte marker GFAP was observed in any of the culture conditions (data not shown). Taken together, these findings show that single overexpression of either OLIG2 or SOX10 in lt-NES cells is not sufficient for efficient OL specification.

We then tested the combined overexpression of both SOX10 and OLIG2 (**Fig. 1a**). Based on our previous findings that undifferentiated lt-NES cells exhibit basal SOX10 expression^5^, we used a fixed concentration of Tet-ON-Olig2-hygro lentivirus (corresponding to multiplicity of infection (MOI) 0.2) and tested two different concentrations of Tet-ON-Sox10-puro, corresponding MOIs of 0.03 and 0.07, respectively. OLIG2 and SOX10 (low and high MOI) were overexpressed in the presence or absence of a cocktail of maturation factors (SAG, PDGFAA, AA, T3, NT3 and IGF-1), extensively used for OL production in previous studies^10, 12^. We first confirmed the expression of both transcription factors in the generated cells by immunocytochemistry (**Fig. 1b**). Interestingly, we found that after 7 days of induction, about 80% of the human lt-NES cells expressed the OL marker O4 without differences in percentage between the different protocols (in terms of presence of maturation factors and SOX10 overexpression levels, for specific percentages for each culture condition, see **Fig. 1c**). Similar percentage of O4+ cells was found in immunocytochemical images at day 10 (**Fig. 1d**). Regarding other neural cell types, no expression of the astrocyte marker GFAP was detected in any of the culture conditions (data not shown) while expression of the neuronal marker TUJ1 was below 5% (**Fig. 1e**).

Taking advantage of the membranous nature of the O4 staining, we could distinguish different cell morphologies within our cultures. This allowed us to subclassify the generated hiOLs into non-ramified, mid-ramified, and highly ramified, corresponding to different stages of maturation (for criteria, see *Methods*). At day 10 of programming, about 3% of the hiOLs generated under different culture conditions exhibited highly ramified morphology except for the condition with high overexpression of SOX10 and the presence of maturation factors, where percentage reached 7.9 ± 3.7% (**Fig. 1f**). A similar trend was found when analyzing the presence of myelin basic protein (MBP)-expressing hiOLs, with about 1.5% for three of the culture conditions and 5.9 ± 1.8% for high overexpression of SOX10 with maturation factors (**Fig. 1g**). All MBP-positive cells exhibited highly ramified morphology. Additionally, the myelin protein PLP1, as well as the mature OL marker CC1 were found at day 10 in all four culture conditions (**Fig. 1h**), suggesting the presence of mature OLs at this time-point.

Given the minimal differences in the numbers of O4+ cells generated across conditions, we decided to use the simplest protocol, consisting in the lower SOX10 overexpression and absence of maturation factors, in all subsequent experiments.

### Transcriptomic analysis reveals a heterogeneous population of hiOLs at different stages of maturation

To further characterize the molecular identity of the generated OLs, we compared their transcriptomic profile, using single nucleus RNA sequencing, with those from adult and fetal human cortical brain cells obtained from neurosurgery and medical abortions, respectively (**Fig. 2a, b**). We first verified the existence of a population of cells co-expressing OLIG2 and SOX10 (**Fig. 2c, d**) as well as the OL marker O4 (**Fig. 2e, f**), which constituted 8.1% and 25.9% of the total number of cells, respectively, for adult and fetal cortical tissue as analyzed by flow cytometry (**Fig. 2g, h**).

**Fig 2.**
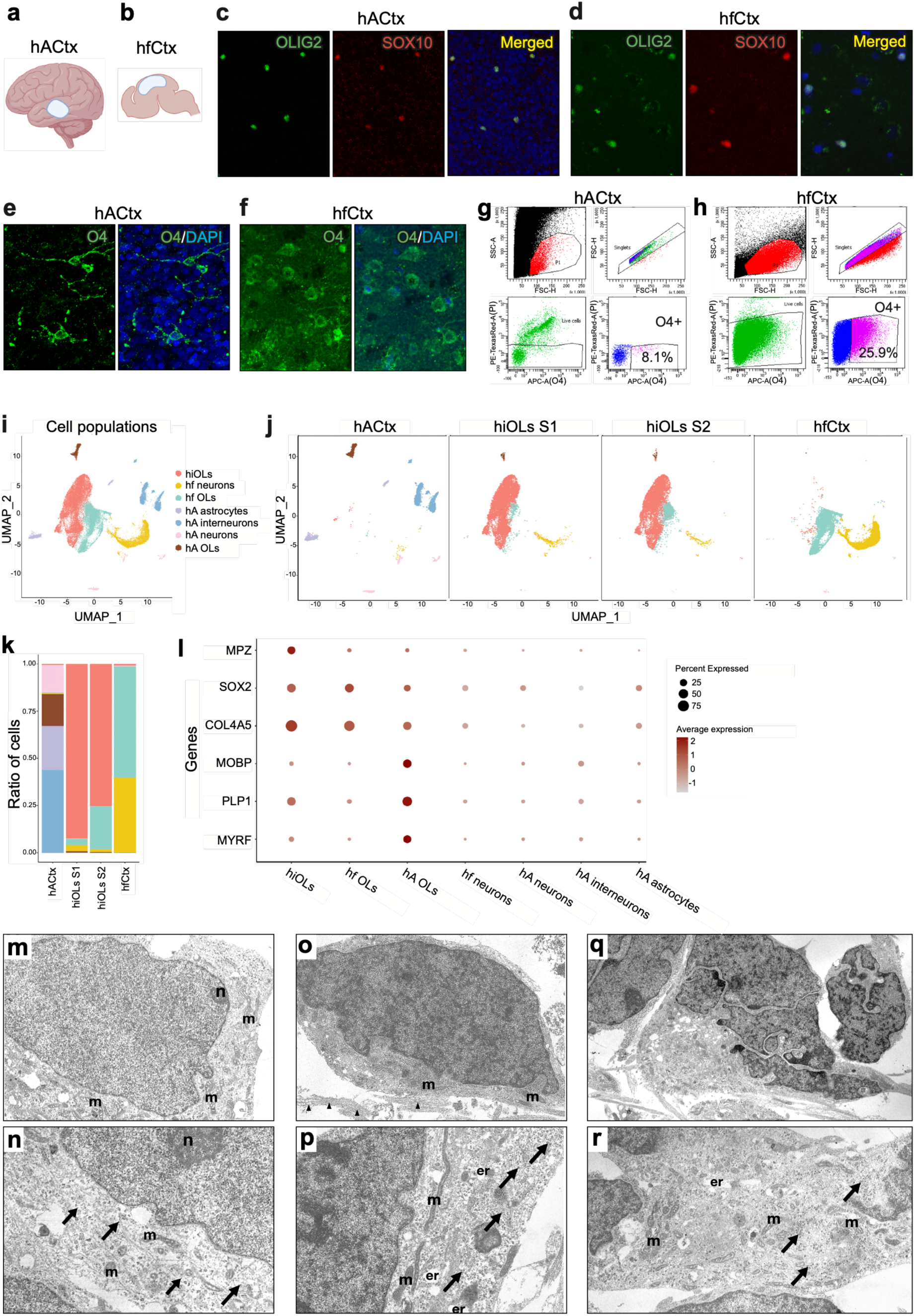
Transcriptomic and ultrastructural characterization of human induced oligodendrocytes (hiOLs) *in vitro*. **a**, **b** Image of the location where human (**a**) adult and (**b**) fetal tissue was obtained. **c**-**f** Confocal images showing expression of the transcription factors OLIG2 and SOX10 (**c**, **d**) and the OL marker O4 (**e**, **f**) in human adult cortical tissue. **g**, **h** Dot plots showing the percentage of the O4+ cells, OL marker, in human adult (**g**) and fetal (**h**) cortical tissue. **i** UMAP plot annotated with the identified cell types. **j** UMAP plot showing cell types obtained from all four samples. **k** Bar plot showing the proportion of different cell types in all four samples. **l** A dotplot showing OL canonical markers in the different clusters. (M-R) Ultrastructural characterization of immature hiOLs in culture. **m** Electron micrograph of an immature OL cell showing a round, electron-lucent nucleus containing predominantly non-condensed euchromatin and a single small nucleolus. **n** An electron-lucent cytoplasm of an immature OL containing small, elongated or oval mitochondria and abundant free ribosomes (arrows). Magnification: ×2900. **o**, **p** Fine structure of human pre-myelinating hiOLs. (**o**) Pre-myelinating OL exhibiting a relatively small volume of moderately electron-lucent cytoplasm, with the nucleus occupying most of the cell body. Several long, thin processes enriched in microtubules are visible (arrowheads). (**p**) Cytoplasm of a pre-myelinating OL containing numerous polyribosomes (arrows), elongated mitochondria, and short, mildly dilated profiles of the endoplasmic reticulum. Magnification: (o) ×3600; (p) ×4800. **q**, **r** Ultrastructural features of human mature hiOLs. (**q**) Mature OL exhibiting an electron-dense appearance and an irregularly shaped nucleus with prominent chromatin clumping along the inner nuclear membrane. (**r**) Electron-dense cytoplasm of a mature OL containing short cisternae of rough endoplasmic reticulum, short mitochondria, and polyribosomes (arrows). Magnification: (q) ×1900; (r) ×2900. Abbreviations: m – mitochondria; er – endoplasmic reticulum; n – nucleolus. Images in A and B were created using Biorender.

After sequencing and quality control we obtained a dataset of 40182 nuclei from 4 samples (2 independent hiOL productions containing a pool of cells at day 7 and 14 of induction, one fetal tissue and one adult human cortex, **Supplementary Fig. 2a-d**). Single-cell-nuclei transcriptomic analysis revealed distinct cell clusters identified in both fetal and adult human samples, corresponding to neurons, OLs, and astrocytes (**Fig. 2i, j**). Among the hiOLs samples, 0.3-0.6% of the cells clustered with adult OLs, 3.4-22.7% with fetal OLs, 1.5-3.4% with fetal neurons, and 0.1-0.2% with other minor populations. Notably, a majority (75.3-92.5%) of the hiOLs formed a cluster with a unique transcriptomic profile (**Fig. 2k**). This cluster lacked expression of neuronal-(*GAD2*, *BCL11B*, *GRIN1* and *RBFOX3*) (**Supplementary Fig. 2f, g**) and astrocyte-(*ALDH1L1*, *GFAP* and *AQP4*) (**Supplementary Fig. 2f, h**) related genes. Moreover, expression of undifferentiated neuroepithelial stem cell–associated genes (*HES5*, *PAX6* and *SOX1*) was low or absent (**Supplementary Fig. 2f, i**). Importantly, we found expression of the genes *MPZ*, *MOBP*, *PLP1* and *MYRF*, present in the adult OLs cluster, and expression of the genes *SOX2* and *COL4A5*, present in both fetal and adult OLs, in hiOLs (**Fig. 2l** and **Supplementary Fig. 2e**). These results showed that induction of SOX10 and OLIG2 in human lt-NES cells generates a heterogeneous population of OL-like cells sharing genes from both fetal and adult human OLs.

### Ultrastructural analysis confirms the presence of hiOLs in different stages of maturation

We next analyzed the ultrastructural characteristics of the hiOLs using transmission electron microscopy (TEM). TEM analysis of the cell cultures at day 14 post-induction revealed a population of lineage-specific OLs at distinct maturation stages, with no evidence of contaminating neuronal or astrocytic phenotypes. Within the cultures, we identified a population of cells exhibiting ultrastructural characteristics consistent with immature OLs (**Fig. 2m, r**). These cells displayed round-to-oval, electron-lucent nuclei containing predominantly non-condensed euchromatin (**Fig. 2m**). A single small nucleolus was frequently observed, along with a thin peripheral rim of heterochromatin. The electron-lucent cytoplasm contained short, mildly dilated cisternae of the endoplasmic reticulum, small, elongated or oval mitochondria, and abundant free ribosomes, consistent with a metabolically active and immature cellular state (**Fig. 2n**)^17^. In addition, a distinct population corresponding to pre-myelinating OLs (pre-OLs) was observed (**Fig. 2o, p**). These cells were characterized by a relatively small volume of moderately electron-lucent cytoplasm, with the nucleus occupying most of the cell body (**Fig. 2o**). Nuclear chromatin was moderately condensed and more compact than that observed in OPCs, reflecting progression toward differentiation. The cytoplasm of pre-OLs contained numerous polyribosomes, elongated mitochondria, and short, mildly dilated profiles of the endoplasmic reticulum (**Fig. 2p**). Pre-OLs extended several long, thin processes enriched in microtubules, indicating increased cytoskeletal organization and early morphological maturation; microtubules were generally oriented parallel to the long axis of the processes (**Fig. 2o**)^18^. A third population identified in the cultures corresponded to mature OLs based on their ultrastructural features (**Fig. 2q, r**). These cells exhibited an overall electron-dense appearance, with irregularly shaped nuclei showing prominent chromatin clumping along the inner nuclear membrane (**Fig. 2q**). The cytoplasm of mature OLs was similarly electron-dense and contained short cisternae of rough endoplasmic reticulum, short mitochondria, and polyribosomes, consistent with a fully differentiated OL phenotype (**Fig. 2r**)^19^.

To determine the relative abundance of these populations, we performed a semi-quantitative morphometric analysis based on the classification of cells sampled from randomly acquired, non-overlapping TEM fields across independent cultures. This analysis showed that the majority of hiOLs has the intracellular molecular features of immature OLs (83 ± 6%), whereas smaller fractions were classified as pre-myelinating OLs (15 ± 5%) and mature OLs (2 ± 1%), consistent with the immunocytochemical analysis and transcriptomic profile of hiOLs.

Together, these findings indicate that combined OLIG2 and SOX10 overexpression efficiently drives oligodendroglial lineage specification.

### Generation of hiOLs via OLIG2 and SOX10 overexpression is robust and reproducible

Next, we confirmed that our results were not specific to a single cell line. We repeated the induction process in two additional human neuroepithelial cell lines, one male, NES-12-7, and one female, NES-C001H (**Supplementary Fig. 3a, b**). Consistent with the results above, we found 79.6 ± 4.8% and 56.6 ± 8.1% of O4+ cells at day 7 of induction, respectively (**Supplementary Fig. 3c, d**). Moreover, the percentages of O4+ cells at day 10 were 82.1 ± 13.2% and 80.1 ± 7.1% (NES-12-7 and NES-C001H, respectively, **Supplementary Fig. 3e**) with a similar proportion of non-ramified and ramified cells and the presence of MBP+ cells in the cultures (**Supplementary Fig. 3f-h**)

To assess reproducibility, the described protocol was executed by four independent researchers in our lab using the same human lt-NES cell line. As a result, the percentage of O4+ cells at day 7 of induction was 86.9% with a standard deviation of 4.1%.

We also evaluated whether the generated OLs could be cryopreserved. hiOLs at day 7 of induction were frozen and thawed 3 months later. The percentage of O4+ cells at day 10 was 76.0 ± 5.2% (**Supplementary Fig. 4a**). The percentage of non-ramified was 75.7 ± 1.6%, mid-ramified 18.0 ± 1.7%, and highly ramified 4.5 ± 2.1% (**Supplementary Fig. 4b**). These results were consistent with freshly generated OLs.

Finally, we examined whether by modifying the culturing conditions from attachment to suspension we could affect the proliferation ratio or the level of maturation of the generated cell product. For that, hiOLs were generated via OLIG2 and SOX10 overexpression under low-attachment conditions (**Supplementary Fig. 5a, c, d**). In this format, cultures formed free-floating spheres (**Supplementary Fig. 5b**). At day 9 post-induction, the proportion of O4+ cells reached 72.5 ± 3.8%, comparable to attached cultures, as determined by flow cytometry and confirmed by immunocytochemistry (**Supplementary Fig. 5e, f**). Spheres were disaggregated and fixed 24 h later (10 DIV). OL morphology based on O4 staining was similar to hiOL generation in attached cultures (**Supplementary Fig. 5g**). Moreover, mature OLs were also found as shown by expression of MBP and CC1 in the spheres and the presence of highly ramified O4+ cells post-disaggregation (**Supplementary Fig. 5h-j**). We found no differences in percentage of proliferative cells (assessed by Ki67 expression, **Supplementary Fig. 5l, m**) with 10.1 ± 0.9% in high attachment protocol versus 14.2 ± 4.2% in low attachment conditions, indicating that in both culture conditions the OLs preserve a degree of mitotic activity. Differences were found in the percentage of cells in a more immature stage, as judged by NESTIN expression, with a significantly higher percentage under low-attachment conditions (43.7 ± 2.8% versus 13.6 ± 1.4%, **Supplementary Fig. 5n, o)**.

### Human induced OLs myelinate human axons both *in vitro* and after allotransplantation into adult human cortical tissue

To assess the functionality of the generated hiOLs, we cultured hiOLs on day 7 of induction on 3D cell culture surfaces containing nanofibers. After 2 weeks, about 15% of the total number of O4-expressing hiOLs showed a highly ramified morphology (**Fig. 3a**). They also exhibited myelination capacity, as evidenced by the ensheathment of nanofibers (**Fig. 3b, c**), the expression of MBP, and the presence of the paranodal marker Caspr (**Fig. 3d, e**). Ensheathment of nanofibers was also evident when hiOLs were cultured under low attachment conditions (**Supplementary Fig. 5k**).

**Fig 3.**
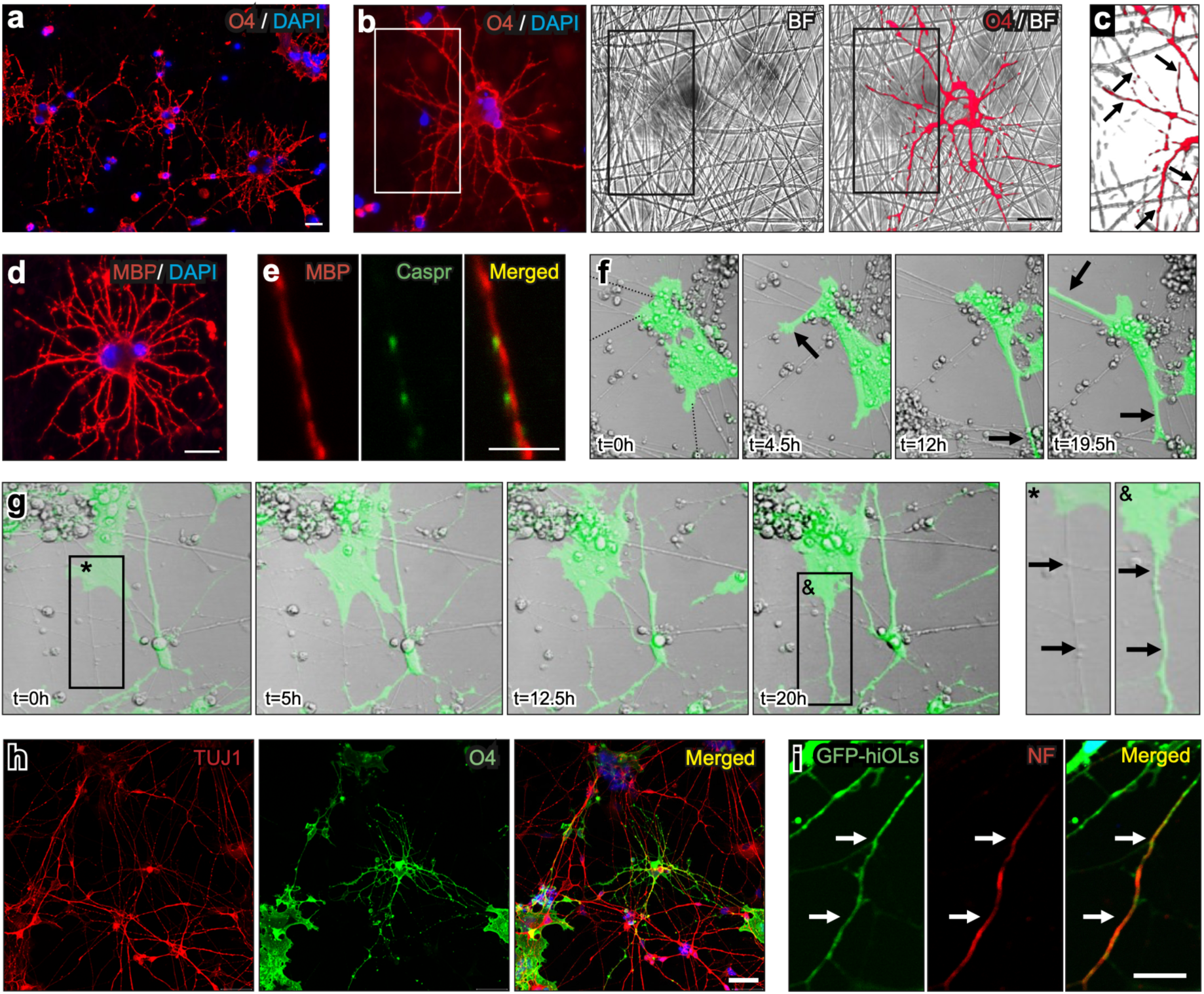
Human induced oligodendrocytes (hiOLs) myelinate nanofibers and human axons *in vitro*. **a** Representative confocal image of O4+ cells after 2 weeks in 3D cell culture surfaces containing nanofibers. **b** Representative image of a O4+ cell in close contact with nanofibers. **c** High amplification image showing colocalization of hiOL processes and nanofibers. **d** hiOL showing expression of myelin basic protein (MBP) after 2 weeks in contact with nanofibers. **e** Colocalizations of a MBP projection and the paranodal marker Caspr. **f** Images extracted from live confocal imaging in cocultures of GFP-labeled hiOLs and human lt-NES induced neurons. Time-lapse of dynamic morphological changes is shown. Arrows show hiOLs processes targeting axons. **g** Time lapse of a hiOL enwrapping an axon in 20 h (* unwrapped axon and & wrapped axon). **h** Representative immunofluorescence image of a co-culture of hiOLs (stained with O4) with lt-NES cells-induced neurons (stained with TUJ1) at day 21 of programming (day 14 of co-culture). **i** The arrows show hiOL processes in close proximity to the neurofilament (NF) marker. Scale bars in a, b, d, i, 20 μm. Scale bar in e, 5 μm.

Further evidence for axonal ensheathment *in vitro* was evaluated in co-cultures of GFP-tagged hiOLs and human lt-NES-induced neurons (lt-NES-iNs, generated via overexpression of NGN2^20^). To visualize the dynamic morphological changes of hiOLs in the presence of neurons, we performed live imaging from days 12 to 14 of induction. The video recordings showed high motility of hiOLs in culture, extending processes towards multiple neuronal projections (**Fig. 3f**; **Supplementary Video 1**). Some initially non-myelinated axons were ensheathed by hiOLs processes in less than 24 h, as shown by time-lapse imaging and by colocalization of OL processes and axonal marker (**Fig. 3g-i; Supplementary Video 2**).

Since grafted OLs in a potential therapeutic setting must be able to interact with diverse neuronal populations in the brain, we conducted co-cultures of GFP-hiOLs with both excitatory neurons (**Supplementary Fig. 6a, b**), lt-NES-iNs, and inhibitory neurons, generated from ES cells through small molecule differentiation (**Supplementary Fig. 6c, d**)^21^. After 14 days in co-culture, we detected areas in which OL processes were aligned with neuronal projections from both excitatory and inhibitory neurons (**Supplementary Fig. 6b, d**).

Moving towards a more clinically relevant setting, we assessed the capacity of the generated hiOLs to survive, differentiate and myelinate in an adult human *ex vivo* system where architecture and distribution of different cell populations are conserved^22^. The GFP-labeled hiOLs programmed for 7 days were transplanted onto adult human cortical organotypic slices and characterized 4 weeks after grafting (**Fig. 4a**). We identified graft-derived GFP+ cells distributed throughout the tissue, extending beyond the transplantation site, and exhibiting complex morphology and arborization (**Fig. 4b**). Overexpression of OLIG2 and SOX10 was maintained for 1 week after transplantation, when doxycycline was removed from the media. We first confirmed that the transplanted cells retained their OL identity after induction was stopped, as shown by the expression of OLIG2 and SOX10 (**Fig. 4c**). Moreover, colocalization of GFP+ cells with the mature OL marker CC1 were found (**Fig. 4d**), arguing for the presence of mature graft-derived OLs. Importantly, no colocalizations with neuronal marker NeuN or astrocytic marker STEM123 (human GFAP) with grafted cells were found (**Fig. 4e**), indicating that the graft was composed exclusively by OL lineage cells.

**Fig 4.**
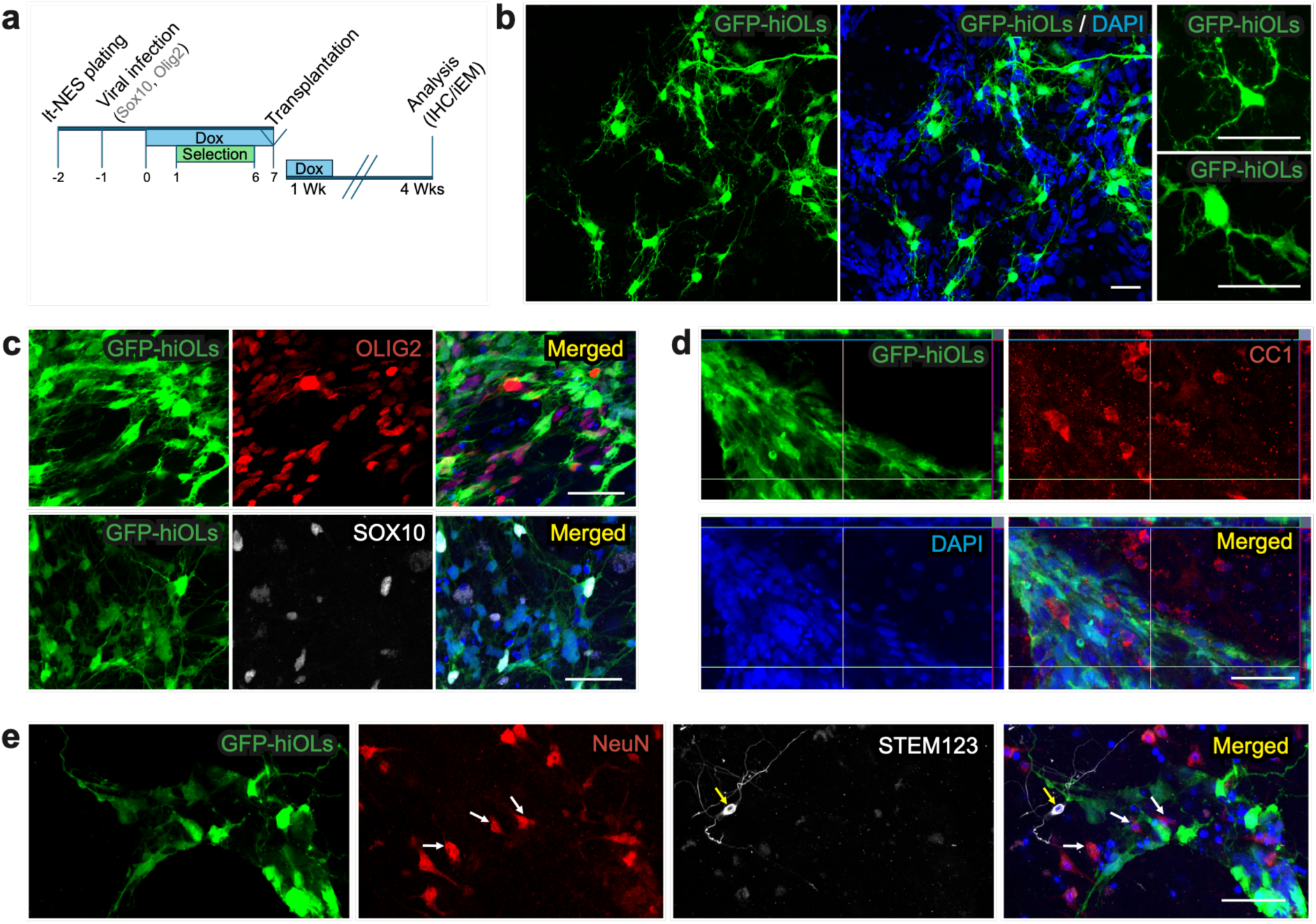
Generation of oligodendrocyte (OL)-based grafts following allotransplantation of OLIG2- and SOX10-overexpressing lt-NES cells into organotypic adult human brain cultures. **a** Schema of the allotransplantation of GFP-labelled human induced OLs into human adult cortical tissue. Panel created in PowerPoint Version 16.107.4. **b** Confocal image showing an overview of the graft 4 weeks after transplantation. **c** Representative confocal images showing the expression of OLIG2 and SOX10 in grafted hiOLs. **d** Orthogonal confocal images of hiOLs expressing the mature OL marker CC1, 4 weeks post-transplantation. **e** Grafted hiOLs show lack of colocalization with neuronal (NeuN, white arrows) and astrocytic (human GFAP, also known as STEM123, yellow arrow) markers. Nuclear staining (DAPI, blue) is included in the merged panel. Scale bars, 50 μm.

To validate the immunohistochemical results, we performed ultrastructural characterization of the grafted cells. One month post-transplantation, immunoelectron microscopy (iEM) revealed that GFP+ immunogold-labeled cells exhibited the ultrastructural hallmarks of mature, myelinating OLs, with notable heterogeneity in nuclear and cytoplasmic electron density (**Fig. 5a**). These cells displayed dense cytoplasm and nuclei with irregular chromatin distribution, including prominent clusters of heterochromatin abutting the inner nuclear membrane (**Fig. 5a,b**). The cytoplasm contained a granular endoplasmic reticulum (ER) composed of short, dilated cisternae, occasionally organized into parallel stacks (**Fig. 5b**). Mitochondria exhibited characteristic ultrastructural features, and free polysomes were abundant, while glycogen granules were infrequent. The Golgi apparatus was well developed, and electron-dense bodies with either homogeneous or heterogeneous content were present. While microfilaments were sparse, microtubules were abundant and predominantly aligned longitudinally within OL processes (**Fig. 5c**). Immunogold particles were diffusely distributed throughout the cytoplasm, with exclusion from mitochondria and the ER (**Fig. 5a-c**). Collectively, these features suggest the identity of the cells as mature myelinating OLs.

**Fig. 5.**
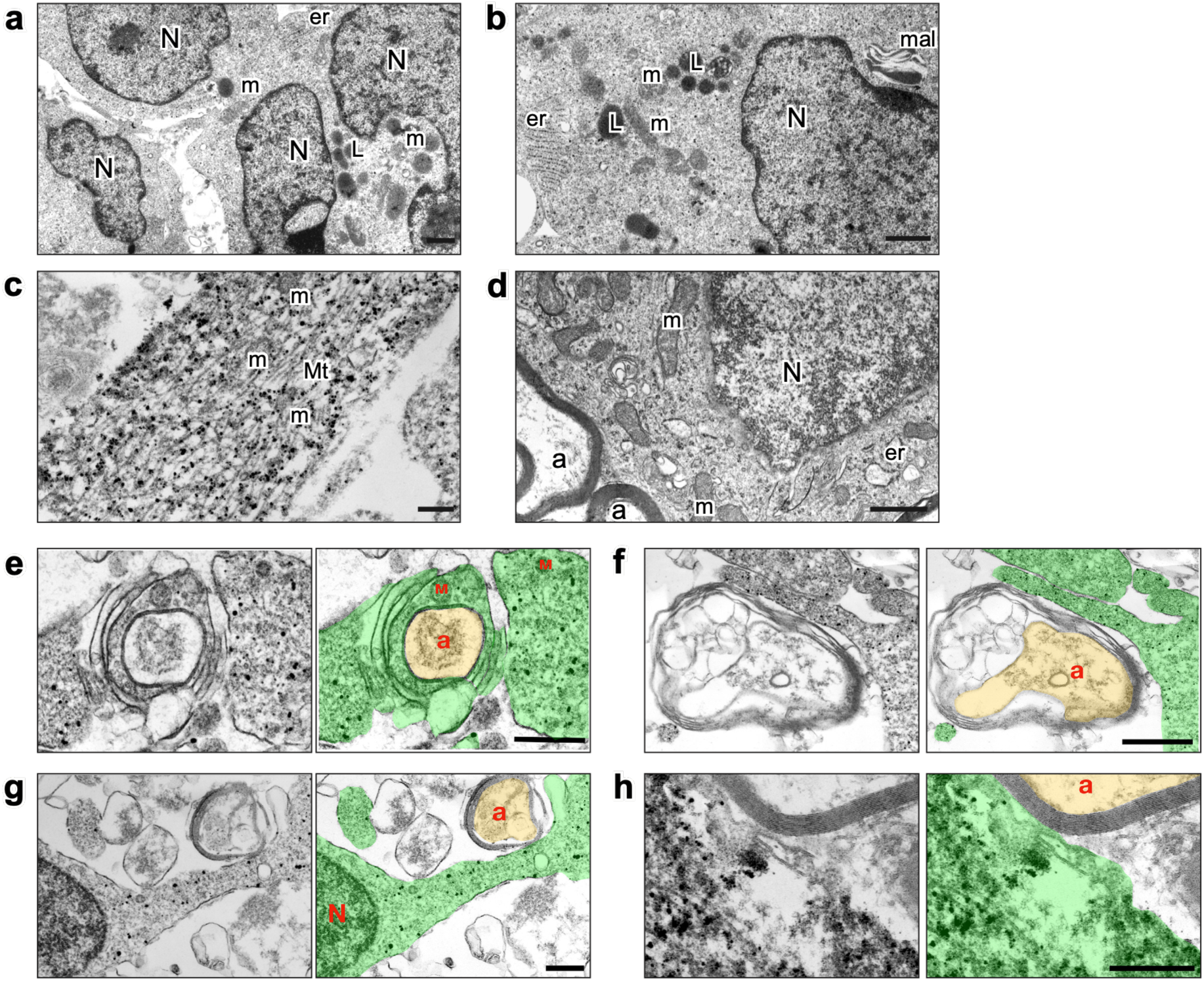
Ultrastructural characteristics of human induced oligodendrocytes (hiOLs) grafted into adult human brain tissue. **a-d** Transmission electron micrographs showing the morphology of mature, GFP+ immunogold-labeled hiOLs derived from human lt-NES cells four weeks following transplantation into adult human cortical slices. **a** GFP+ cell exhibiting mature OL morphology. **b** Nucleus with uneven chromatin distribution; granular ER with stacked cisternae; mitochondria with characteristic morphology; presence of myelin-associated lysosomes. **c** Abundant, longitudinally aligned microtubules within OL processes. **d** Endogenous mature OL in the human cortex. Abbreviations: N – nucleus, m – mitochondria, mt – microtubules, er – endoplasmic reticulum, a – myelinated axon, l – lysosome, mal – myelin-associated lysosome. Scale bars in a, b, d = 1 µm; c = 0.5 µm. **e-h** Transmission electron micrographs showing stages of myelination of host axons by GFP+ hiOL-derived processes. **e** Early myelination. **f**, **g** Intermediate myelination. **h** Compact myelin. For clarity, GFP+ hiOL-derived processes and myelin are pseudocolored green; host axons are pseudocolored yellow. Abbreviations: N – nucleus, m – mitochondria, a – myelinated axon. Scale bars in e–g, 1 µm; h, 0.5 µm.

Interestingly, a subset of OLs contained lysosomes exhibiting lamellar or membrane-like structures (**Fig. 5b**), reminiscent of myelin-associated lysosomes, which have been implicated in myelin biogenesis and may play a role in sorting and packaging myelin proteins.

All observed ultrastructural features in these graft-derived OLs were indistinguishable from those of endogenous human cortical OLs (**Fig. 5d**).

Importantly, GFP+ OL-derived processes formed myelin sheaths around host axons, as evidenced by numerous immunogold-labeled processes adjacent to or enveloping unlabeled host axons (**Fig. 5e-h**). The ensheathment displayed classical features, including inner and outer cytoplasmic tongues in close apposition to the developing or compact myelin. Axons at varying stages of myelination were identified, ranging from initial wrapping to complete ensheathment with compact myelin (**Fig. 5e-h**).

## DISCUSSION

Our demonstration that cells generated efficiently and rapidly by our programming protocol produce grafts composed exclusively of OLs myelinating human axons provides the basis of a new cell therapeutic strategy in demyelinating disorders. We conclude that the generated human cells are *bona fide* functional OLs based on several lines of evidence. First, they express key OL markers, including O4, the myelin proteins MBP and PLP1, and the mature OL marker CC1, as shown by flow cytometry, single cell sequencing and immunocytochemistry. Second, they exhibit ultrastructural features of OL in different stages of differentiation *in vitro*, as revealed by TEM. Third, they ensheathe axons when co-cultured with human neurons derived from pluripotent stem cells. Finally, after transplantation into organotypic slices of adult human brain tissue, the cells retain a stable OL phenotype, confirmed by marker expression and ultrastructural characteristics of mature OLs observed by iEM. Importantly, graft-derived myelin sheets are found around the adult human host axons.

Several strategies have previously been developed with the aim of generating human OLs with therapeutic potential from either pluripotent stem cells or fibroblasts. These protocols are robust and not dependent on a specific cell line^9–12, 23^. In addition, two studies have demonstrated that the generated OLs can be cryopreserved^9, 11^, as shown for our cells, an essential property for a therapeutic product. While protocols based on the addition of small molecules are usually time-consuming, taking from 55 to 100 days^7, 8, 24^, ectopic expression of different transcription factors implicated in OL specification accelerates the process. The induction of only SOX10 or a combination of three transcription factors (SOX10, OLIG2, NKX6.2) in human iPSC-derived NPCs gives rise to 50% of O4+ OLs in 10 days and 70% of O4+ OLs in 28 days, respectively^10, 11^. Here, using human lt-NES cells, we produce 80% of O4+ OLs within 7 days of overexpression of SOX10 and OLIG2, demonstrating the fastest and most efficient protocol for the generation of human OLs described to date. A recent study used a protocol similar to ours on human iPS cells (also based on the overexpression of OLIG2 and SOX10) and obtained comparable percentages of induction, confirming the reproducibility of our protocol. The myelination capacity of the generated oligodendrocytes was not reported^25^.

The lt-NES cells used here exhibit the advantages of pluripotent stem cells, such as stability and long-term expandability, but, importantly, lack tumorigenic potential^26, 27^. Moreover, previous protocols based on small molecule differentiation or reprogramming strategies have relied on the addition of a specific cocktail of growth factors and small molecules (including SAG, PDGF, AA, T3, NT3 and IGF-1, among others) to direct the differentiation into OLs^8, 10–12^. We found that this cocktail was not needed for efficient hiOL production using our programming protocol in human lt-NES cells, reducing both protocol complexity and production costs. Moreover, in contrast to conventional protocols requiring extracellular matrix substrates, we have shown that using low-attachment culture conditions, the efficiency of hiOL generation is not affected, which reduces variability, simplifies handling, and enhances compatibility of the protocol with scalable and GMP-compliant manufacturing processes.

It is an advantage that the present protocol can produce virtually pure OL grafts. We previously showed that human iPS cell-derived lt-NES cells, subjected to a cortical priming protocol, gave rise to grafts containing both neurons and OLs^5^. Similarly, previously published protocols have generated not only OLs but also neurons or astrocytes^9, 10^. While such mixed grafts may be valuable for conditions involving loss of both neurons and OLs, such as ischemic stroke, they are not optimal for treating demyelinating disorders, in which the therapeutic goal is to remyelinate axons before neuronal degeneration occurs. Moreover, most demyelinating disorders, including leukodystrophies and multiple sclerosis, primarily affect white-matter regions^28, 29^ with minimal neuronal presence, where the introduction of inappropriate cell types could disrupt local circuitry. One solution has been to sort the OL lineage cells prior to transplantation. However, sorting procedures can compromise cell viability and trigger stress-related intracellular responses^30^. Moreover, despite sorting, neurons and astrocytes were still detected in the graft^10^.

Cell-based strategies based on human OL transplantation have shown promise for regenerating the CNS and contributing to functional recovery in preclinical models of hypomyelination or demyelination^7, 23, 31–33^ or spinal cord injury^34^. However, only a few clinical trials exploring the therapeutic use of intracerebral transplantation of human stem cell-derived products aiming at axonal remyelination have so far been performed. Intraspinal transplantation of human ES cell-derived OL progenitor cells was evaluated in Phase I and Phase I/IIa clinical trials in five and twenty-five individuals with spinal cord injury, respectively. The results demonstrated long-term safety over ten years with no tumor formation, neurological decline, or immune rejection after immunosuppression withdrawal. Cases of modest sensory and/or motor improvements were described^35–37^. Similar results were found in Phases I to II clinical trials where transplantation of autologous olfactory ensheathing cells, HuCNS-SC (NPCs derived from fetal CNS tissue), NSI-566 (a human fetal spinal cord-derived NSC) or Schwann cells (obtained from sural nerve biopsies) was performed in patients with SCI with follow-up to 3-6 years^38^. All these cell sources have demonstrated remyelination capacity in preclinical studies; however, whether this occurs in patients or whether signs of recovery arise from other mechanisms is unknown. A clinical trial with allogenic transplantation of human neural stem cells was performed in four patients with central hypomyelination in Pelizaeus-Merzbacher’s disease^39, 40^. After 2 to 5 years, patients exhibited diffusion MRI changes at the transplantation site and in distant brain areas, suggesting increased myelination. Finally, two clinical trials in patients with multiple sclerosis using intrathecally transplanted human fetal NPCs reported reduced brain atrophy at the highest dose and increased cerebrospinal fluid levels of anti-inflammatory and neuroprotective molecules. The beneficial effects were attributed to bystander mechanisms rather than to graft-derived axonal remyelination^41, 42^.

A major challenge in moving cell therapies toward clinical application for patients with neurological diseases is that nearly all preclinical studies are conducted in xenograft settings (human cells transplanted into animal brains). While rodents and humans share neural cell types, key differences exist in gene expression, synaptic plasticity, neuronal networks, and responses to physiological and pathological stimuli. To ensure the development of safe and efficient cell therapies, preclinical models that enable analysis of human cells within human brain tissue are urgently needed (Wechsler et al. 2018). Importantly, the human OLs generated by our protocol survive and function in the adult human brain tissue environment, strongly supporting their feasibility for clinical translation. Previous clinical attempts using human OL transplantation have been based on xenotransplantation experiments and translating results from animal models to humans are known to present significant challenges. Our findings support the idea that effective stem cell-based treatments for myelin repair will become possible in several human diseases. In demyelinating disorders with a strong autoimmune component, such as multiple sclerosis, OL transplantation most likely must be complemented with treatments to suppress neuroinflammation^43, 44^ or genetic modification of the transplanted cells^45^ to resist the disease process. Further investigations will explore these strategies to determine the therapeutic potential of our hiOLs for treating demyelinating disorders.

## METHODS

### Human iPS cell-derived lt-NES cell culture

Generation and maintenance of human iPS cell-derived lt-NES cells were performed as described previously^16, 22^. Briefly, lt-NES cells were cultured in flasks pre-coated with poly-L-ornithine (1:100, Cat# P3655, Merck Millipore) and mouse laminin (0.002 mg/mL, Cat# 23017015, ThermoFisher Scientific) in the presence of a basic medium composed of DMEM-F12 (Cat# 11320074, ThermoFisher Scientific), N2 supplement (1:100, Cat# 17502001, ThermoFisher Scientific), and glucose (1.6 g/L, Cat# G8769, Sigma-Aldrich) supplemented with growth factors: basic fibroblast growth factor (bFGF) (Cat# AF-100-18B, Peprotech), epidermal growth factor (EGF) (Cat# AF-100-15, Peprotech) and B27 (Cat# 17504001, ThermoFisher Scientific), all of them at the final concentration of 10 ng/mL. Cells were fed every day with growth factors and passaged every third to fourth day using 0.025% trypsin (Cat# T4049, Sigma), followed by trypsin inhibitor (0.5 mg/mL, Cat# 17075029, ThermoFisher Scientific).

### Vector cloning and virus production

For generating the Tet-ON-Sox10-Puro, we first digested the TetO-Ngn2-Puro (Cat# 52047, Addgene) with EcoRI and XbaI to remove the Ngn2 factor. We next amplified the Sox10 factor from the Tet-O-FUW-Sox10 (Cat# 45843, Addgene) using the forward primer 5’-GAATTCGCCACCATGGCCGAG-3’, which contains the EcoRI restriction site followed by the Kozak sequence and the 5’ region of the Sox10 coding sequence, and the reverse primer 5’-TCTAGAAGGTCGGGATAGAGTCGTATATAC-3’, which contains the 3’ region of the Sox10 coding sequence without the stop codon followed by the XbaI restriction site. We then TOPO-cloned the PCR product and digested it with EcoRI and XbaI, to finally clone it into the opened tetO-Puro backbone obtained in the first step. For the generation of the Tet-ON-Olig2-Hygro, we used the tetO.Nfib.Hygro plasmid (Cat# 117271, Addgene) and the service from Genscript to replace the Nfib coding sequence with the Olig2 coding sequence.

Lentiviral vectors were produced according to^46^ and were in titers of 10^7^-10^8^ TU/mL as determined by qRT-PCR.

### Generation of hiOLs in attachment culture conditions

The lt-NES cells were transduced with three lentiviral vectors: Tet-ON-Sox10-Puro, Tet-ON-Olig2-Hygro and FUW-M2rtTA (Tet-controlled transactivator. Cat# 20342, Addgene). The expression of both transcription factors, SOX10 and OLIG2, was initiated at day 0 by administration of 2.5 µg/mL doxycycline (dox) (Cat# D9891, Sigma-Aldrich) and kept until day 10. On day 1, basic medium was replaced with glial differentiation medium (GM) composed of DMEM-F12, N2 supplement (1:200), B27 supplement without retinoic acid (1:100, Cat# 12587001, ThermoFisher Scientific) and 1% Glutamax (0.25 µg/mL, Cat# 35050061, ThermoFisher Scientific). Antibiotic selection was carried out on days 1 and 3 for puromycin (1.25 µg/mL, Cat# A11138-03, ThermoFisher Scientific), and from days 1 to 5 for hygromycin (0.2 mg/mL, Cat# 10687010, ThermoFisher Scientific). In addition, for the optimization of the protocol, in some culture conditions media were supplemented with smoothened agonist (SAG) (1 µM, Cat# S7779, Selleckchem), ascorbic acid (AA) (200 µM, Cat# A4544, Sigma), insulin-like growth factor 1 (IGF-1) (10 ng/mL, Cat# 100-11, Peprotech), neurotrophin-3 (NT3) (10 ng/mL, Cat# 450-03, Peprotech), platelet-derived growth factor (PDGF) (10 ng/mL, Cat# 100-13A, Peprotech) and triiodothyronine (T3) (20 ng/mL, Cat# T6397, Sigma-Aldrich).

On day 7, cells were detached for characterization and destined to either flow cytometry analysis or further differentiation using immunocytochemistry.

### Generation of hiOLs in 3D spheres

The same protocol as described for attachment conditions was followed up to day 3 of induction. At this time-point, cells were detached and seeded at a density of 1 × 10^6^ cells on a 6-well plate with ultra-low attachment surface (Cat# 3471, Corning). Dox was added every day, and half of the glial medium was refreshed every other day until day 9, when spheres were either dissociated for flow cytometry and posterior seeding on coverslips or collected for immunocytochemistry.

### *In vitro* characterization of hiOLs by flow cytometry

The hiOLs were harvested on day 7, washed and resuspended in 200 µL of antibody master-mix: Anti-O4-APC antibody (Cat# 130-119-155, Miltenyi) diluted 1:200 in FACS buffer (PBS + 2% fetal bovine serum + 2 nM ethylenediaminetetraacetic (EDTA)). Cells were incubated for 45 min at +4° C in darkness and washed with FACS buffer by centrifugation. Finally, cell pellets were resuspended in 300 µL of propidium iodide (PI) diluted in FACS buffer (1:1000) at least 5 min before acquisition.

The flow cytometry analysis was performed on a BD LSR Fortessa-X20 flow cytometer. The data were analyzed using BD FACSDiva 9.0 software (BD Biosciences) and FlowJo v.10.8 Software (BD Life Sciences). As for the gating strategy, viable single cells were selected from the total events by size, complexity, and PI negativity.

### *In vitro* characterization of hiOLs by immunocytochemistry

The hiOLs, plated on coated glass coverslips, were fixed at day 10 of the programming protocol in 4% paraformaldehyde (PFA) for 10 min at room temperature. Cells followed a first step of permeabilization and blocking using KPBS-0.025% Triton X-100 (ThermoFisher Scientific) with 5% of normal donkey serum (NDS) (Cat# S30-M, Merck Millipore) for 45 min at room temperature. After that, primary antibodies diluted in blocking solution were applied overnight at +4 °C, followed by fluorophore-conjugated secondary antibodies diluted in blocking solution for 1 h at room temperature. Nuclei were then stained for 10 min at room temperature with 4′,6-diamidino-2-phenylindole (DAPI) diluted 1:1000 in KPBS. Stained glass coverslips were mounted on slides using a mounting media containing DABCO. For the stainings requiring antigen retrieval, an initial incubation with sodium citrate, pH 6.0, Tween 0.05% at +65°C for 30 min was performed.

The following primary antibodies were used: mouse anti-CC1 (1:200, Cat# ab16794, Abcam), goat anti-DCX (1:400, Cat# SC11365, Santa Cruz), rabbit anti-GABA (1:2000, Cat# A2052, Sigma), mouse anti-GFAP (STEM123, 1:1000, Cat# Y40420, Takara), chicken anti-GFP (1:1000, Cat# AB16901, Merck Millipore), rabbit anti-KGA (1:1000, Cat# ab9343, Abcam), rabbit anti-ki67 (1:500, Cat# ab16667, Abcam), chicken anti-MAP2 (1:1000, Cat# ab5392, Abcam), mouse anti-MBP (1:500, Cat# 808401, BioLegend), mouse anti-Neurofilament Axonal Pan (1:1000, Cat# SMI-312R/837904, BioLegend), mouse anti-Neurofilament Neuronal Pan (1:1000, Cat# SMI-311R/837801, BioLegend), mouse anti-O4 (1:100, Cat# MAB1326, R&D Systems), rabbit anti-Olig2 (1:500, Cat# ab109186, Abcam), rabbit anti-PLP1 (1:200, Cat# ab254363, Abcam), goat anti-Sox10 (1:400, discontinued, Santa Cruz), rabbit anti-Sox10 (1:500, Cat# ab180862, Abcam), rabbit anti-TUJ1 (1:500, Cat# 802001, BioLegend) and mouse anti-TUJ1 (1:500, Cat# 801202, BioLegend).

For quantification of the percentage of cells positive for the OL markers O4 and MBP, 10-µm-thick Z-stack images were acquired with a 20X objective using a confocal microscope (LSM 780, Zeiss, Germany). A total of 10 images spaced 1000 µm apart were taken per coverslip per condition. Maximum intensity projection images were analyzed using ImageJ software. The percentage of positive cells for each marker was quantified as follows: marker+ cells / total DAPI+ cells. Cell counting was performed independently by three blinded researchers, with the final result representing the average value.

For the morphological analysis of the hiOLs, a classification was established based on morphological complexity according to the number of processes using the cytoplasmic staining O4: unbranched cells were classified as “non-ramified”; cells with less than 5 ramifications were termed “mid-ramified” and cells with more than 5 ramifications were classified as “highly ramified”.

### Human adult and fetal brain samples

The research on human tissue was conducted in accordance with the highest ethical standards and in agreement with Swedish legislation. Written consent was obtained and the participants were informed about the purpose of the research.

Human fetal forebrain tissue was obtained from material available after elective termination of pregnancy at the University Hospital in Malmö, Sweden, in accordance with guidelines approved by the Lund/Malmö Ethical Committee (Dnr 6.1.8-2887/2017).

Human adult neocortical tissue was obtained with informed consent by resection of a small piece of the middle temporal gyrus from patients undergoing elective surgery for temporal lobe epilepsy, in accordance with guidelines approved by the Regional Ethical Committee, Lund (Dnr. 2021-07006-01).

### Nuclei isolation from fetal and adult brain tissue

The nuclei isolation from adult and fetal cortical tissue was performed as previously described^47^ using a sucrose gradient-based isolation. For the human fetal tissue, cells were disaggregated prior to nuclei isolation. For the human adult tissue, due to the decreased cell viability post-disaggregation, tissue was snap-frozen after collection in surgery room and a small section of the tissue (3 mm^3^) was cut from the original sample tissue and dissociated in 1 ml of ice-cold lysis buffer (0.32 M sucrose, 5 mM CaCl_2_, 3 mM MgAc, 0.1 mM Na_2_EDTA, 10 mM Tris-HCl, pH 8.0, 1 mM DTT) using a 1 ml tissue douncer (Wheaton). The lysate was then carefully layered on top of a 2.8 ml sucrose cushion (1.8 M sucrose, 3 mM MgAc, 10 mM Tris-HCl pH 8.0, and 1 mM DTT diluted in milliQ water) and then centrifuged at 30,000 × g for 2 hours and 15 min. Once the supernatant was removed, the pelleted nuclei were softened for 10 min in 50 μl of nuclear storage buffer (15% sucrose, 10 mM Tris-HCl pH 7.2, 70 mM KCl, and 2 mM MgCl_2_, all diluted in milliQ water), resuspended in 300 μl of dilution buffer (10 mM Tris-HCl, pH 7.2, 70 mM KCl, and 2 mM MgCl_2_, diluted in milliQ water) and then filtered through a cell strainer (70 μm). The nuclei were sorted via fluorescence-activated nuclear sorting (FANS) with a FACS Aria (BD Biosciences) at +4 °C at low flow rate using a 100 μm nozzle (reanalysis showing >95% purity).

A detailed protocol can be found in^48^.

#### Single-nucleus RNA sequencing

The nuclei for single nuclei RNA sequencing (30000 nuclei per sample) were centrifuged at 500g 10 min at +4 °C. The supernatant was carefully removed so that the remaining volume ended up being 37 ul. The nuclei were suspended by flicking the tube and pipetting gently up and down a few times and then loaded onto the Chromium GEM-X 3’ Chip along with the reverse transcription master mix according to the manufacturer’s protocol for the Chromium GEM-X single cell 3’ kit v4 (10X Genomics ref 1000711) to generate single-cell gel beads in emulsion. cDNA amplification was achieved following the guidelines from 10X Genomics, employing 13 cycles of amplification of the 3’ libraries. Sequencing libraries were generated with unique dual indices (TT set A) and pooled for sequencing on a Novaseq X plus using a 100-cycle kit and 28-10-10-90 reads.

### Single nuclei RNA Sequencing Analysis

To obtain count matrix files sample-specific FastQ reads were aligned to the human genome assembly (version GRCh38) using the Cell Ranger pipeline (‘cellranger count’ 10x Genomics Cell Ranger 9.0.0) with default parameters and include-introns set to true.

All downstream analyses were performed using R (v4.3.1)2 and the standard Seurat (v4.4.0)3 workflow^49^. Low-quality nuclei and genes were filtered out based on the fraction of the total number of genes detected in each nuclei (± 3 nmads). Nuclei with gene counts between 600 and 10,000 were retained for further downstream analysis. Additionally, nuclei with less than 3% mitochondrial transcripts were included for the analysis (percent.mt <3). All four samples were merged using the Harmony algorithm (‘sample’)^50^. Any doublets or multiplets detected using scDbIFinder were removed.

For visualization and clustering, manifolds were calculated using UMAP methods (RunUMAP, Seurat) and 20 precomputed principal components and the shared nearest neighbor algorithm modularity optimization-based clustering algorithm (FindClusters, Seurat) with a resolution of 0.1. This resulted in 12 clusters, which were annotated using canonical known markers gene expression for respective cell types.

### Transmission electron microscopy (TEM) of hiOLs culture

The hiOL cultures were fixed with 2% formaldehyde and 0.2% glutaraldehyde in 0.1 M phosphate buffer (pH 7.4). After fixation, samples were washed with deionized water and post-fixed in 1% osmium tetroxide in 0.1 M phosphate buffer for 30 minutes on ice. Dehydration was performed through a graded ethanol series, followed by infiltration with ethanol/epoxy resin mixtures. Samples were then embedded in pure epoxy resin and polymerized at 60 °C for 48 hours. Ultrathin sections were cut, stained with lead citrate, and examined using a JEM-100CX transmission electron microscope (JEOL, Japan).

### Morphometric quantification of OL lineage cells by TEM

For quantitative assessment of oligodendroglial populations, a morphometric analysis was conducted on TEM images acquired at day 14 post-induction.

Ultrathin sections were systematically imaged across the grid for each well using a predefined sampling strategy. Non-overlapping fields of view were acquired in a randomized manner to ensure representative coverage of each culture and to minimize sampling bias. Fields were selected based solely on section quality and the absence of technical artifacts, without consideration of cellular composition.

Only cells with a clearly identifiable nucleus and sufficient surrounding cytoplasm to allow reliable ultrastructural classification were included in the analysis. Cells that were partially sectioned, morphologically ambiguous, or affected by sectioning or staining artifacts were excluded.

All images were anonymized and randomized prior to analysis. Cells were assigned to one of three categories – immature OLs, pre-myelinating OLs, or mature OLs – based on predefined ultrastructural criteria, including nuclear chromatin organization, cytoplasmic electron density, organelle composition, and the presence of microtubule-containing cellular processes.

For each well, at least 100 cells were counted manually across multiple fields of view. The proportion of each cell type was calculated relative to the total number of classified cells per well. Data are presented as mean ± SD, where variability reflects differences across four independent wells.

### Co-culture of hiOLs with human neurons and hiOL culture in nanofibers

For the generation of excitatory neurons, human iPS cell-derived lt-NES cells were subjected to dox-induced overexpression of neurogenin-2 (NGN2) followed by antibiotic (puromycin) selection for 3 days as described previously^51^. Human inhibitory neurons were generated from ES cells as described^21^.

For the establishment of neuron-OL cultures, hiOLs programmed for 7 days were plated on top of coverslips containing either excitatory or inhibitory mature neurons (ratio 10,000 OLs/100,000 neurons) in the presence of neuronal induction medium (DMEM/F12 supplemented with N2 [1:100, Thermo Fisher Scientific, Sweden] and B27 [1:50, Thermo Fisher Scientific, Sweden]). Dox was daily added to the media until day 9 and every other day from day 9 to day 14.

Co-cultures were either fixed after 14 days in coculture for characterization via immunocytochemistry or subjected to live-cell imaging from day 12 to day 14.

For the 3D culture experiment of myelination, NanoECM® plates (Neuromics), containing randomized nanofiber polymers, were pre-coated with poly-L-ornithine and laminin. hiOLs at day 7 of induction were reseeded at a density of 50,000 cells per chamber in glial medium supplemented with dox. Half of the medium was changed every other day, and cells were fixed in 4% PFA after 14 days for immunocytochemistry analysis.

### Live cell imaging

The GFP+hiOLs and lt-NES-iNs were transferred to the incubation chamber of a Zeiss LSM780 laser scanning confocal microscope. The cultures were maintained and imaged for 2 days at +32 °C in 5% CO2, with fresh medium added daily.

Fluorescence and brightfield time-lapse images were acquired every 30 min using a 20X dry (air) objective. Time-lapse single-plane images were collected to monitor the samples over time. The 488 nm argon laser was used to detect GFP expression in GFP+ cells, with the signal collected using a Gallium Arsenide Phosphide (GaAsP) detector. The same 488 nm argon laser was employed for brightfield imaging of non-labeled neuronal cells, with the transmitted light captured using a Transmitted Photomultiplier Tube (T-PMT). Differential Interference Contrast (DIC) imaging was utilized to enhance the contrast of the transmitted light images, facilitating detailed visualization of the sample’s structural features. The Zeiss LSM780 was operated using Zen Black software version 2012.

Time-lapse videos and individual frames extracted from these videos, in which GFP+hiOLs had migrated toward neuronal cells and overlapped with their axonal structures, were analyzed. Particular attention was paid to detecting overlap between the GFP signal and axonal structures, as this phenomenon strongly indicates interactions between OLs and neurons and possibly early myelination events.

### Organotypic cultures of human adult cortical tissue

The tissue was collected and handled as described previously^22^. Briefly, the resected tissue was immediately placed in ice-cold modified human artificial cerebrospinal fluid and sectioned into 300 μm slices using a vibratome (Leica VT200S). Slices were kept in a rinsing medium (HBSS [1X, Cat# 14175095, Thermo Fisher Scientific], HEPES [4,76 g/L, Cat# A1069, AppliChem], Glucose [2 g/L, Cat# G7021, Sigma-Aldrich] and Penicillin/Streptomycin [500 U/mL, Cat# 15140122, Thermo Fisher Scientific]) before they were either fixed with 4% PFA for the acute characterization of the tissue or transferred to cell culture inserts (Cat# PICMORG50, Millipore, scaffold membranes) for long-term culture. Organotypic culture was performed in human adult cortical medium (Brain Phys medium [without phenol red, Cat# 05791, StemCell Technologies] supplemented with B27 [1:50], glutamax [1:200], gentamicin [50 mg/mL, Cat# 15750037, Thermo Fisher Scientific], Antibiotic/Antimycotic [1X, Cat# 15240062, Thermo Fisher Scientific], BDNF [50 ng/mL, Cat# 450-02, Peprotech] and NT3 [50 ng/mL, Cat# 450-03, Peprotech]), in 5% CO2 at +37 °C.

### Transplantation of hiOLs onto human organotypic cortical slices

Human slices were maintained in culture for 4 days before transplantation of GFP+ hiOLs. The GFP+ hiOLs were harvested at day 7 of induction. After trypsinization, cells were resuspended to a concentration of 100,000 cells/μL in pure cold Matrigel Matrix (Corning), collected into a cold glass capillary and injected as small drops stabbing the semi-dry slice at various sites. After Matrigel was solidified, additional medium was carefully added to immerse the tissue. The media was changed every other day. Dox was added until day 14 of programming (1 week after transplantation). Tissue was fixed 4 weeks post-transplantation for graft characterization via immunohistochemistry and immuno-electron microscopy (iEM).

### Immuno-electron microscopy (iEM) in human organotypic cortical slices

For pre-embedding immunogold labeling, samples were blocked and permeabilized in PBS containing 2% NDS and 0.05% Triton X-100 for 30 min. They were then incubated overnight at +4 °C with a primary goat anti-GFP antibody (1:150, Abcam) diluted in PBS with 2% NDS and 0.05% Triton X-100, followed by PBS washes. A secondary Nanogold®-Fab’ rabbit anti-goat antibody (1.4 nm gold-conjugated, 1:200, Nanoprobes) was applied for 2 h, and samples were washed again. Fixation was performed with 2% glutaraldehyde in PBS for 30 min, followed by washing with deionized water. Silver enhancement was carried out using the HQ kit (Nanoprobes) under darkroom conditions for 8 min. After enhancement, samples were washed with deionized water and post-fixed in 0.2% osmium tetroxide in 0.1 M phosphate buffer for 30 min on ice. They were then stained with 0.25% uranyl acetate in 0.1 N acetate buffer for 1 h. Dehydration was performed through a graded ethanol series, followed by infiltration with ethanol/epoxy resin mixtures. Finally, the samples were embedded in epoxy resin and polymerized at +60 °C for 2 days. Ultrathin sections were cut, stained with lead citrate, and examined using a JEM-100CX transmission electron microscope (JEOL, Japan).

### Immunohistochemical analysis of human organotypic cortical slices

For staining in organotypic cultures, slices were fixed with 4% PFA overnight at +4°C followed by overnight incubation at +4 °C in permeabilization solution (0.02% Bovine serum Albumin [BSA], 1% Triton X in PBS). Next, the slices were incubated overnight at +4 °C in blocking solution (KPBS, 0.2% Triton X-100, 1% BSA, Sodium azide [NaN3] [1:10,000] and 10% NDS).

Primary antibodies were diluted in a blocking solution and incubated for 48 h at +4 °C. Secondary antibodies, also diluted in blocking solution, were applied for 48 h at +4 °C. Following this, the slices were washed and stained with DAPI for 2 h at room temperature. Finally, they were rinsed with deionized water and mounted with Dabco mounting media.

The following primary antibodies were used: mouse anti-CC1 (1:200, Cat# ab16794, Abcam), anti-mouse GFAP (STEM123, 1:1000, Cat# Y40420, Takara), chicken anti-GFP (1:1000, Cat# AB16901, Merck Millipore), rabbit anti-NeuN (1:500, Cat# ab104225, Abcam) rabbit anti-Olig2 (1:500, Cat# ab109186, Abcam), goat anti-Sox10 (1:400, discontinued, Santa Cruz) and rabbit anti-Sox10 (1:500, Cat# ab180862, Abcam).

Some stainings required antigen retrieval before the permeabilization/blocking step. In those cases, organotypic cultures were incubated with sodium citrate, pH 6.0, Tween 0.05% for 2 h at +65 °C. Quantification of the percentage of grafted cells positive for OLIG2 and SOX10 was quantified as follows: marker+ cells / total GFP+ cells.

### Statistical analysis

The statistical analysis was performed using GraphPad Prism 10.1.1. Statistical significance was tested with the unpaired two-sided *Student t*-test (for comparison of two groups) or Two-Ways ANOVA followed by Sidák’s multiple comparisons post hoc test (for comparison of the 4 different culture conditions in the protocol optimization). P-values<0.05 were considered statistically significant. Data are mean ± SD.

## AUTHOR CONTRIBUTIONS

S.P.-T. conceptualized the study. R.M-C., O.T., P.R-C., A.M.A.K., D.M-H., E.M., I.H., S.K., A.B., A.A., and S.P.-T. conducted the experiments. J.B., E.S., and A.F. generated and/or provided human material. S.P.-T., O.T., R.M-C., O.L., and Z.K. wrote the manuscript. R.M.-C., O.T., Y.S., L.R., D.R.O., H.A., I.C., G.S., and O.L. were involved in data interpretation. Funding for this study was acquired by S.P.-T., Z.K. All authors reviewed, edited and approved the manuscript.

## ACKNOWLEDGMENTS

This work is supported by grants from Crafoordska Stiftelsen (20250660) and the Åke Wibergs Foundation (M23-0041, M24-0186 and M25-0303) to S.P-T. Swedish Research Council (2020-01669), and Swedish Brain Foundation, FO2025-0078-TK-288 to Z.K. Swedish Research Council (2022-02673 to J.J.), the Swedish Brain Foundation (FO2023-0232), Cancerfonden (222185Pj), Barncancerfonden (PR2023-0099) and the Swedish Government Initiative for Strategic Research Areas (MultiPark & StemTherapy) to J.J.. Swedish Research Council, 2023-02409, Hedlund Foundation and Swedish Brain Foundation, FO2022-0132 to HA. Swedish Society for Medical Research (SSMF, S20-0003) to IC. IC is supported by the University of Zurich Research Priority Program ITINERARE – Innovative therapies in rare diseases. The authors acknowledge the Lund Stem Cell Center’s Imaging, FACS and Cell and Gene Therapy Core for the technical support.

## DATA AVAILABILITY

The matrix files and related processed snRNA-seq data from this study are available at (https://tinyurl.com/yeytr4uh). Rest of data are available from the corresponding author on reasonable request.

