## Supplementary information for "Efficient generation of human oligodendrocyte grafts that restore myelin in adult human brain tissue"

[1]Laboratory of Stem Cells and Restorative Neurology, Lund Stem Cell Center, Lund University, 22184 Lund, Sweden; [2]Department of Cytology, Bogomoletz Institute of Physiology; Institute of Genetic and Regenerative Medicine, Strazhesko National Scientific Center of Cardiology, Clinical and Regenerative Medicine, 01024 Kyiv, Ukraine; [3]Laboratory of Molecular Neurogenetics, Department of Experimental Medical Science, Wallenberg Neuroscience Center and Lund Stem Cell Center, BMC A11, Lund University, 221 84 Lund, Sweden; [4]Departamento de Farmacología y Toxicología, Facultad de Medicina, Universidad Complutense de Madrid (UCM), Centro de Investigación Biomédica en Red de Salud Mental, Instituto de Salud Carlos III (CIBERSAM, ISCIII), Instituto de Investigación Hospital 12 de Octubre (i+12), Instituto Universitario de Investigación en Neuroquímica (IUIN), 28040 Madrid, Spain; [5]Neural Stem Cells, Department of Experimental Medical Science, Lund Stem Cell Center, Lund University, Lund, 221 84, Sweden; [6]Stem Cells, Aging and Neurodegeneration group, Department of Experimental Medical Science, Lund Stem Cell Center, Faculty of Medicine, Lund University, 22184, Lund, Sweden; [7]Group of Regenerative Neurophysiology, Lund Stem Cell Center, Department of Experimental Medical Science, Faculty of Medicine, Lund University, 221 84 Lund, Sweden; [8]FACS Core Facility, Lund Stem Cell Center, Lund University, 22184 Lund, Sweden; [9]Division of Metabolism, University Children's Hospital Zurich, University of Zurich, 8008, Zurich, Switzerland; [10]Division of Neurosurgery, Department of Clinical Sciences Lund, University Hospital, 22184 Lund, Sweden.

& Equal contribution

### Supplementary Figures

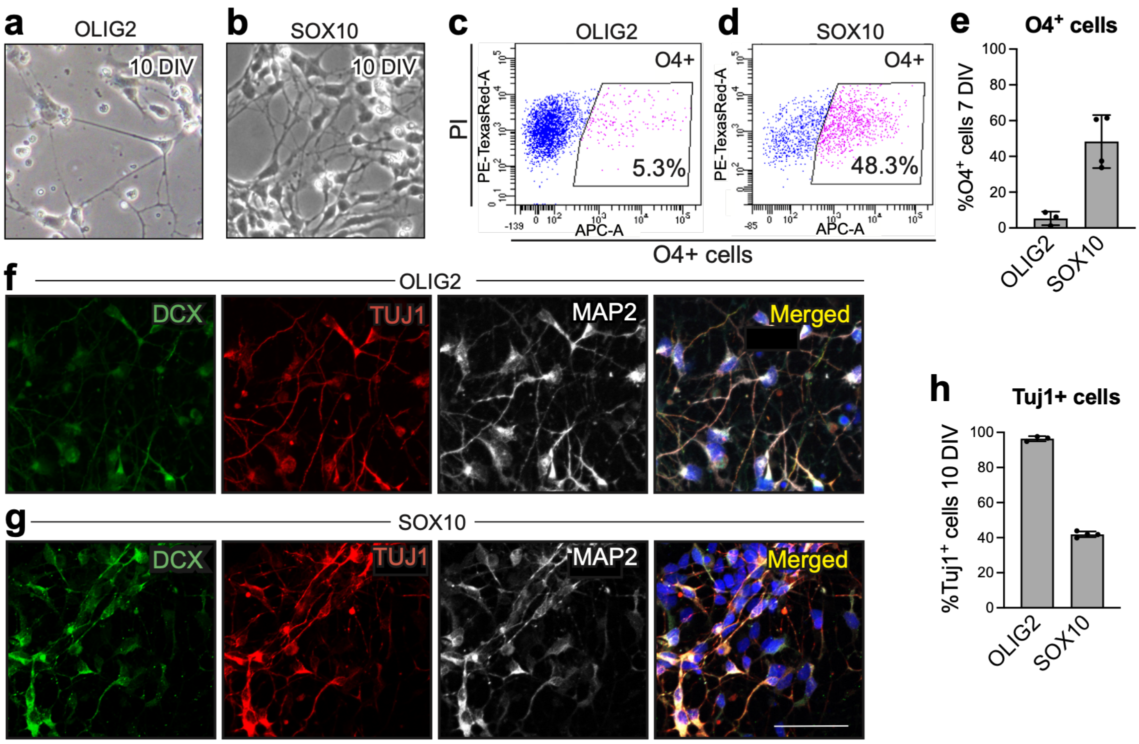


Supplementary Fig. 1. **Human lt-NES cells programmed with either OLIG2 or SOX10 overexpression result in the generation of a mixed population of neurons and oligodendrocytes (OLs). a,b** Brightfield images of human lt-NES cells after 10 days of OLIG2 (**a**) or SOX10 (**b**) overexpression. **c-e** Representative dot plots (**c**,**d**) and quantifications (**e**) of the percentage of cells positive for the OL marker O4 analyzed by flow cytometry at 7 days of programming. **f**,**g** Representative confocal images showing the expression of neuronal progenitor marker doublecortin (DCX), neuronal marker class III beta-tubulin (TUJ1) and the mature neuronal marker microtubule-associated protein 2 (MAP2) at day 10 post-induction. **h** Quantification of cells positive for the neuronal marker TUJ1. Nuclear staining (DAPI, blue) is included in the merged panel. Scale bar, 50 µm. All data are presented as the mean ± SD, n=3-4 biologically independent samples.

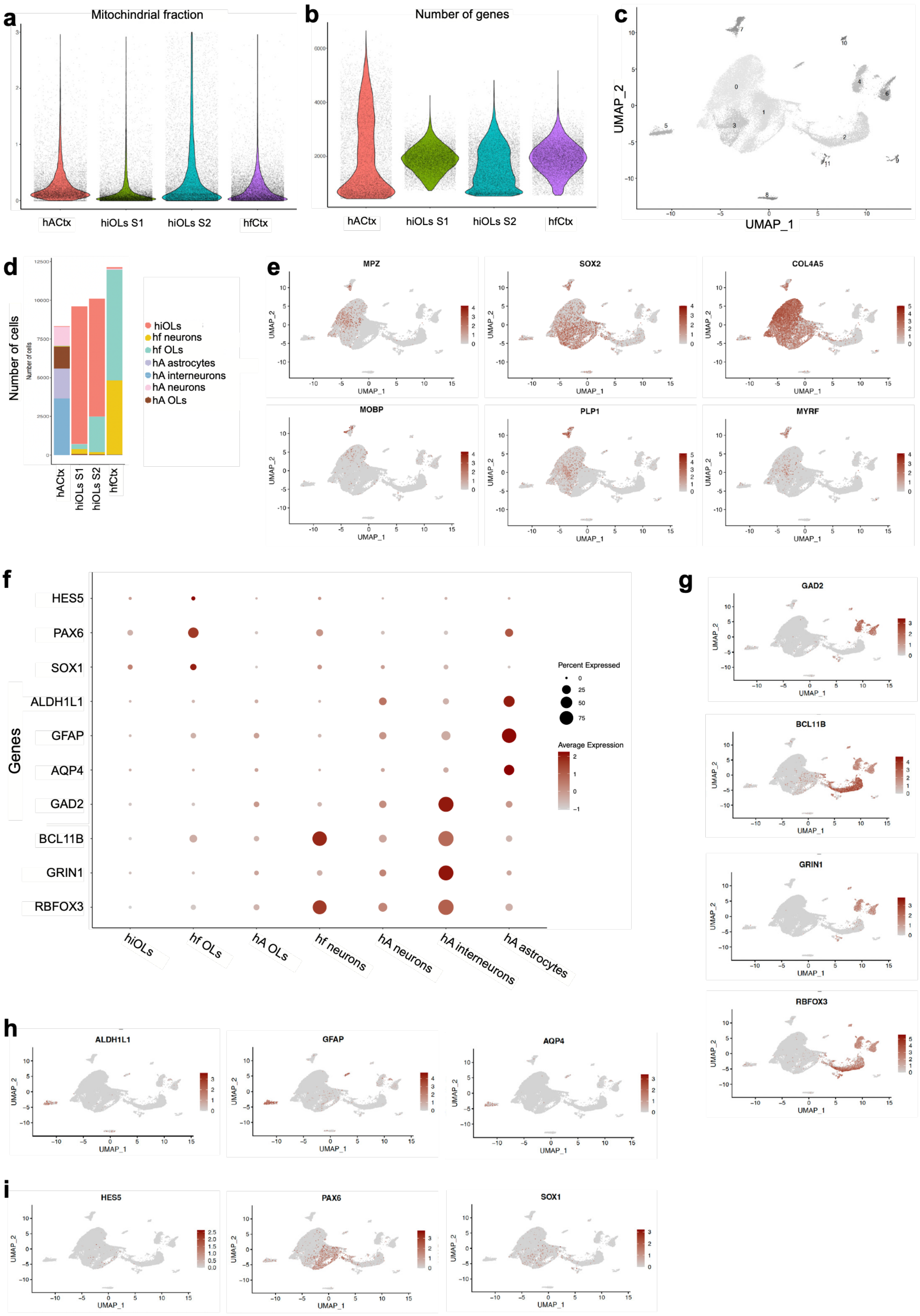


Supplementary Fig. 2. **Transcriptomic profile of human induced oligodendrocytes (hiOLs). a** Violin plot showing the distribution of percentage of mitochondrial genes per nucleus retained for downstream analysis. **b** Violin plot showing number of genes expressed from each nucleus across four samples. **c** UMAP plot annotated with the 12 identified cell clusters. **d** Barplot showing number of nuclei contributed from each celltype across all four samples. **e** Projection of gene expression of selected markers for oligodendrocytes as featureplots. **f** Dotplot showing canonical markers corresponding to respective celltypes. **g-i** Projection of gene expression of selected markers across different celltypes: neurons **(g),** astrocytes **(h)** and lt-NES cells **(i).**

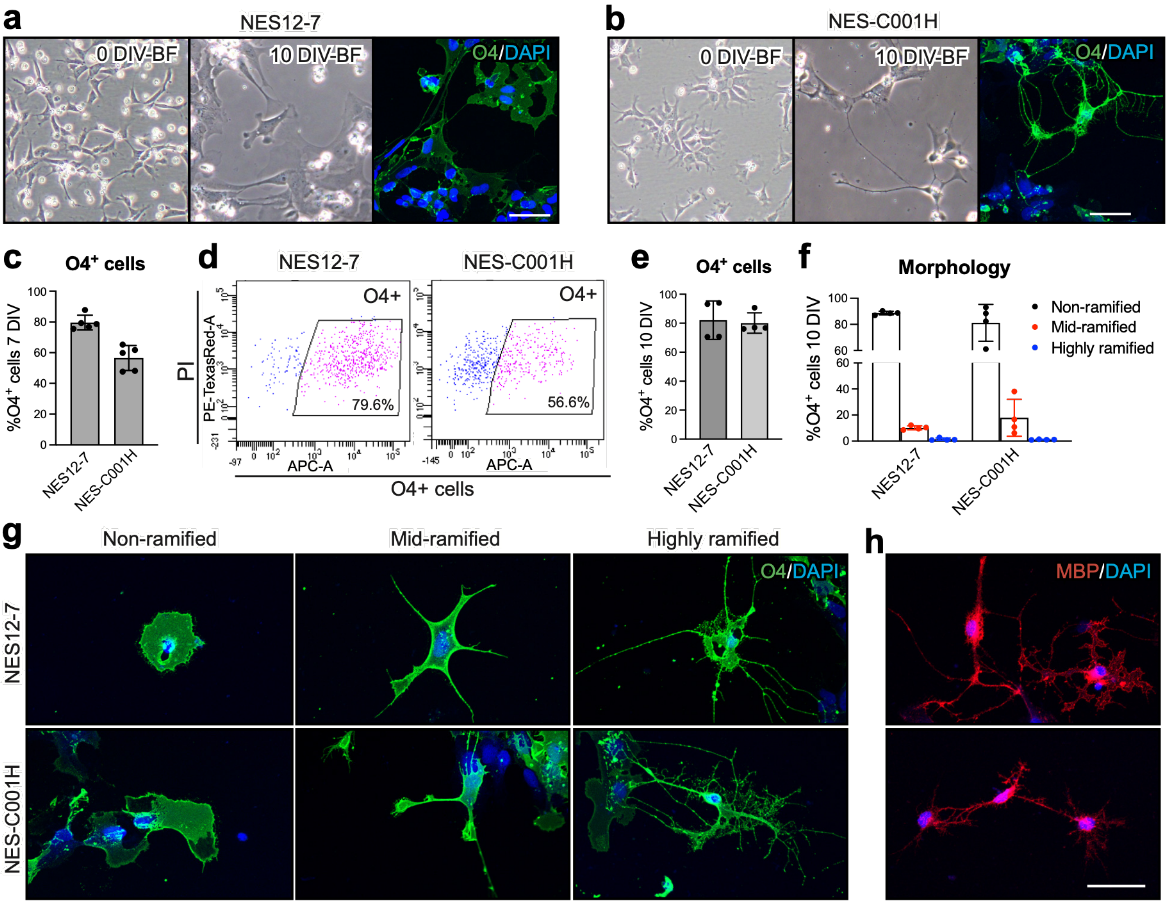


Supplementary Fig. 3. **OLIG2 and SOX10 overexpression in different NES cell lines produce oligodendrocytes (OLs) *in vitro*.** **a-b** Representative brightfield and O4 (OL marker) immunofluorescence images of NES12-7 (**a**) and NES-C001H (**b**) cells at day 0 and 10 of programming. **c,d** Quantification and representative dot plots showing the percentage of the O4+ cells at day 7 of programming. **e** Quantifications of the percentage of O4+ cells analyzed by immunocytochemistry at day 10 of programming. **f** Quantifications of percentages of non-, mid- and highly ramified O4+ cells. **g,h** Representative confocal images of non-, mid- and highly ramified O4+ cells and MBP+ cells. Scale bars in a-b, 50 µm. Scale bars in g-h, 20 µm. All data are presented as the mean ± SD, n=4-5 biologically independent samples.

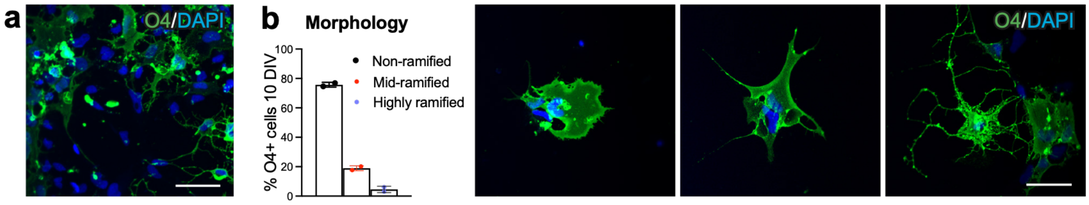


Supplementary Fig. 4. **Cryopreservation of human induced oligodendrocytes (hiOLs). a** Quantification and representative confocal image of O4+ cells 3 days after thawing (10 days of programming). Scale bar, 50 µm. **b** Confocal images and quantification of non-ramified, mid-ramified and highly ramified O4+ cells. Scale bar, 20 µm. All data are presented as the mean ± SD, n=2 biologically independent samples.

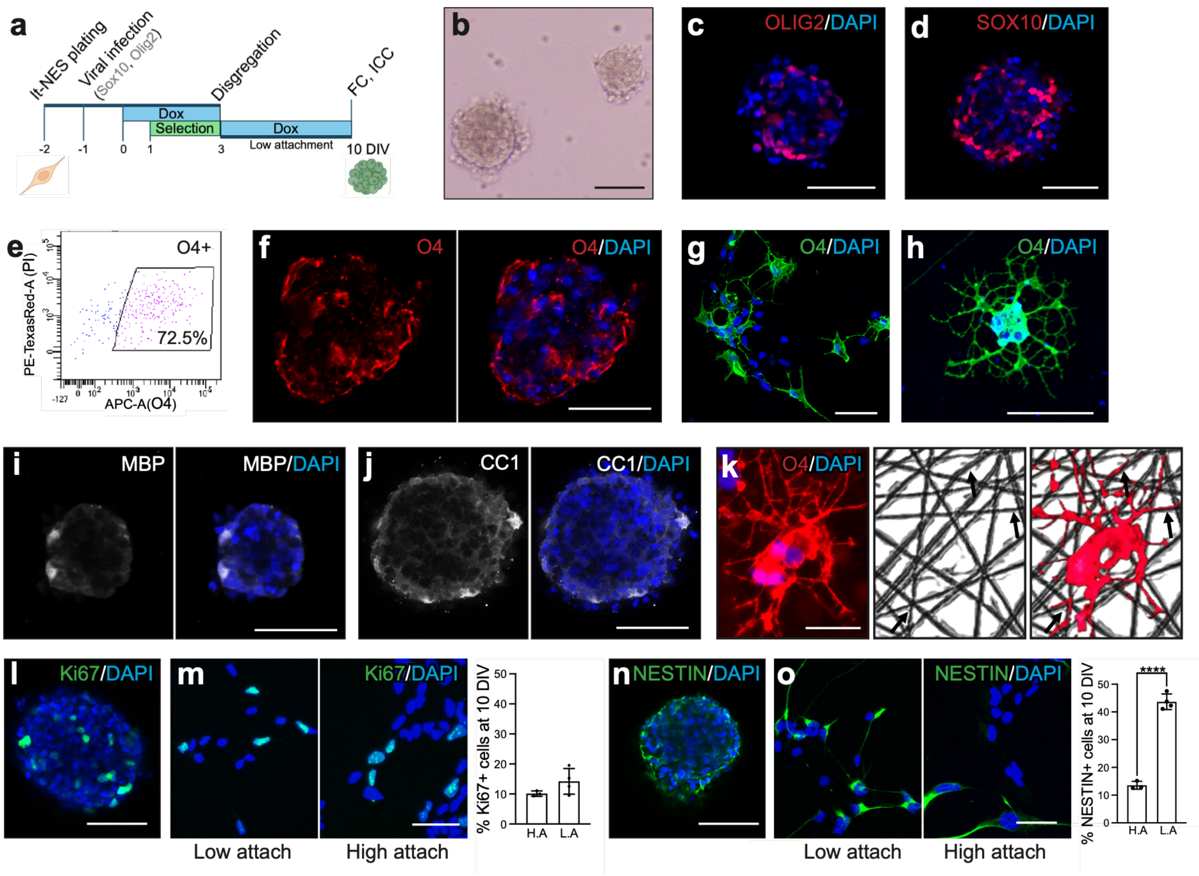


Supplementary Fig. 5. **OLIG2 and SOX10 overexpression in human long-term neuroepithelial-like stem (lt-NES) cells cultured in low attachment conditions efficiently generate oligodendrocytes (OLs) *in vitro* and allows for cell expansion. a** Experimental design to produce human induced OLs from human lt-NES cells using low attachment culture conditions. Dox– doxycycline. FC–Flow cytometry. ICC–Immunocytochemistry. **b** Brightfield image of spheres generated after 9 days of programming in culture. **c, d** Confocal images showing the expression of the transcription factors OLIG2 (**c**) and SOX10 (**d**) **e** Representative dot plot of the percentage of the O4+ cells, OL marker, at day 9 using the low attachment culture conditions. **f, g** Confocal images showing the expression of O4 at day 9 of programming in spheres and 24 h later after disaggregation and culture in attachment conditions. **h** Representative image of a O4+ cell exhibiting highly ramified morphology. **g, i, j** Confocal images showing cells positive for the mature OL markers myelin basic protein (MBP) and CC1 in the spheres. **k** Representative image of O4+ processes in close contact with 3D nanofibers **l, m** Confocal images and quantification of the expression of the proliferation marker Ki67 in hiOLs generated under low and high attachment conditions. **n, o** Confocal images and quantification of the expression of the immature marker NESTIN in hiOLs generated under low and high attachment conditions. All data are presented as the mean ± SD, n=3-4 biologically independent cultures. **a** panel was created in PowerPoint Version 16.107.4. Scale bars in b-j, l and n, 50 µm. Scale bars in k, m and o, 20 µm.

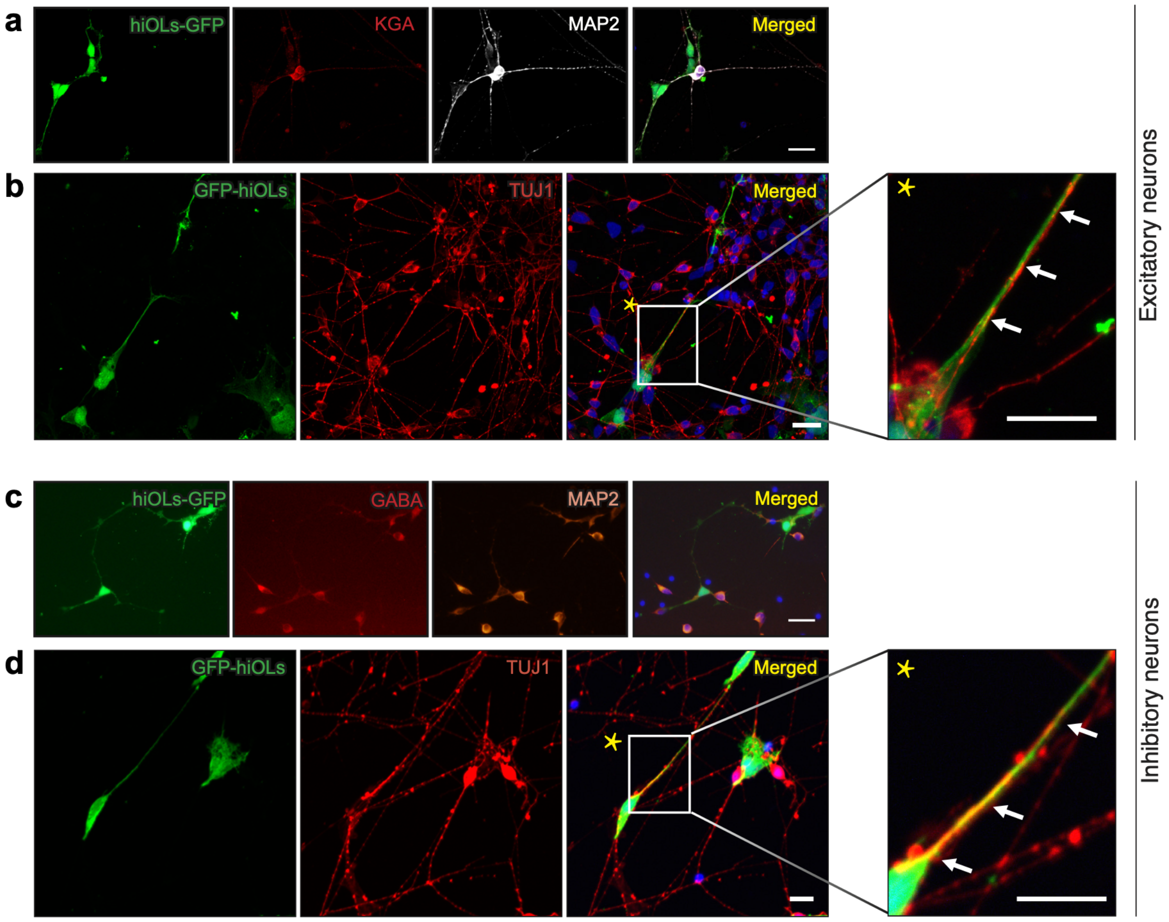


Supplementary Fig. 6. **Human induced oligodendrocytes (hiOL) processes grown in close proximity to excitatory and inhibitory neuronal axons. a** Representative confocal image of the co-culture containing GFP-labelled hiOLs and human lt-NES cells-derived excitatory neurons, stained with glutaminase (KGA) and the mature neuronal marker microtubule-associated protein 2 (MAP2). **b** Image showing excitatory neurons (TUJ1+ cells) and hiOLs (GFP+ cells). **c** Representative confocal image of the co-culture containing GFP-labelled hiOLs and human ES cells-derived inhibitory neurons, stained with gamma aminobutyric acid (GABA) and MAP2. **d** Image showing inhibitory neurons (TUJ1+ cells) and hiOLs (GFP+ cells). Arrows in **b** and **d** show close proximity between both markers. Nuclear staining (DAPI, blue) is included in the merged panel. Scale bars, 20 µm.
